# Biosynthesis of Silver nanoparticles using *Drimia indica* and exploring its antibacterial profiling

**DOI:** 10.1101/2022.07.25.501375

**Authors:** Pratik S. Kamble, Mansingraj S. Nimbalkar, Swaroopa A. Patil

**Affiliations:** Department of Botany, Shivaji University, Kolhapur- 416004

**Keywords:** *Drimia indica*, *Klebsiella*, *Pseudomonas*, *Escherichia*, Microtitre Broth Dilution

## Abstract

Biological synthesis reflects as an eco-friendly, nontoxic and easy method of nanoparticle preparation. Present investigation deals with biosynthesis of silver nanoparticles using *Drimia indica* leaf extract. Initially the synthesized silver nanoparticles were characterized and confirmed by UV-Vis spectroscopy, X-ray diffraction (XRD) and SEM. The synthesized nanoparticles when used for determination of antibacterial activity, by Microtitre Broth Dilution method exhibited remarkable activity against *Pseudomonas aeruginosa, Klebsiella pneumoniae* and *Escherichia coli*.

## 1. INTRODUCTION

Nanotechnology is the most immerging branch of Science and Technology. It is one of the most promising technologies applied in all areas of Science. Apart from its tremendous applications in Pharmaceuticals, textiles industries, electronics etc, it is ecologically sound and cost effective. Metal nanoparticles have received global attention due to their enormous applications in the biomedical and physiochemical fields. Nanoparticles aqueduct the space between bulky materials and molecular structures (Thakkar *et al*., 2010). Synthesizing metal nanoparticles using plants has been recognized as green and efficient way for further exploiting plants as convenient nanofactories (Singh *et. al*., 2016). Nanoparticles with antimicrobial activity are advantageous in reducing acute toxicity, lowering cost and overcoming resistance as compared to other prevalent antibiotics (Pal *et al*., 2007 and Weir *et al*., 2008).

Antibiotics are used as antimicrobial agents in medical field. Higher doses of antibiotics causes harm to human cells and forms cancerous cells and mutations. Use of nanoparticles as antimicrobial agents is increasing day-by-day. The over doses of antibiotics harms human cells, causes paralysis with more disabilities.

Continued use of antibiotics for treatment of an array of infections has imposed the danger of developing antibiotic resistance. Widespread bacterial infection treatment regime involves higher initial doses of antibiotics followed by gradual lowering down the doses, which requires a longer span of time. A scenario of increased initial doses is practiced regularly which has developed antibiotic resistance for a specific microorganism or a set of microorganisms. Treatment plans for dreadful infections having selective antibiotics for use are compromised due to development of antibiotic resistance, as single antibiotic is used for treatment of such infections. Control measures may be possible in case of broad spectrum antibiotic used or in organisms that are non-selective to antibiotics. On the other hand there are infections which need specific/ selective antibiotics for cure. If resistance for single known antibiotic is developed in organisms, the treatment lines may go unattended or may fail totally. There is a need of promising interventions of multifaceted alternative products to antibiotics which can play an vital role in the field of medical science.

Biogenic nanoparticles are used as replacement over antibiotics to target the microbes (Wang *et al*., 2017). Recently, researchers have showed the antimicrobial potential of biogenic nanoparticles on microorganisms. The use of nanoparticles as antimicrobial agents is increasing exceptionally in the field of medical science (Heather., 2017).

Through nanotechnology, highly medicinal plants can be directly used for nanoparticle synthesis. The biologically synthesized nanoparticles have helped in target oriented drug delivery, which has lowered down the drug doses with increased efficacy (Patra *et al*., 2018). The plant based synthesis of nanomaterials can be explored and used for human welfare.

*Drimia indica* (Asparagaceae) is a highly medicinal plant. Is is commonly known as Indian squill or Rankanda or jangali pyaz in India. It is pear-shaped, onion like, scaly bulb that grows up to 30 cm in diameter (Pretorius *et al*., 2005 and Crespo *et al*., 2020). *Drimia indica* contains cardiac glycosides, quinones, resins, saponins and steroids (Chittoor et al., 2012 and Pandey *et al*., 2014). It possesses antiprotozoal, hypoglycemic, anticancer, antidiabetic and antimicrobial properties. (Aswal *et al*., 2019 and Manganyi *et al*., 2021).

## 2. MATERIAL AND METHODS

### 2.1 Materials

The leaves of *Drimia indica* were collected from Dev Dari, Ambheri, District: Satara, State: Maharashtra (GPS– 17.605087°, 74.279927°). Chemicals used were purchased from Thomas Bakers (C) Pvt. Ltd.

### 2.2 Preparation of Plant extract

For the preparation of leaf extract, fresh leaves of *Drimia indica* were washed thoroughly under tap water and bolted dry. Leaves (20 g) were boiled into 100 ml of distilled water at 100 °C for 15 minutes. Extract was passed through Buchner’s funnel and volume of filtrate was adjusted to 100 ml with distilled water.

### 2.3 Synthesis of Silver Nanoparticles

Leaf extract (20%) was taken into burette and 3mM aqueous solution of AgNO_3_ was taken in conical flask. The flask was placed on magnetic stirrer with hotplate at 60°C. Leaf extract was added drop wise into conical flask containing 3mM AgNO_3_ solution with continuous stirring. Colorless AgNO_3_ solution turned yellowish brown which indicated the formation of silver nanoparticles after reduction of silver ions. After 24 hours the reaction mixture was centrifuged at 10000 RPM. The pellet was taken into beaker and washed with alcohol for 2-3 times followed by washes with distilled water. The pellet was dried in microwave oven and further used for characterization process.

### 2.4 Antibacterial Assay

The bacterial cultures were sub cultured on liquid nutrient broth and they were incubated at 37°C for 24 hours in incubator with continuous stirring. The characterized nanoparticles were used for determination of antimicrobial potential by Microtitre Broth Dilution method on spectrophotometer against pathogenic bacteria *Pseudomonas aeruginosa, Klebsiella pneumoniae* and *Escherichia coli*. The antibacterial activity of silver nanoparticles was measured by calculating percent inhibition and minimum inhibitory concentration (MIC).

## 3. Result and Discussion

### UV visible spectroscopy

Synthesis of nanomaterials and their applications are having great importance in modern era. Silver nanoparticles were formed by gradual reduction of silver ions during reaction with *Drimia indica* plant extract. The reaction mixture showed the color change from colorless to light brown to dark brown colored solution.

The initial formation of silver nanoparticles was determined by UV-visible spectroscopy. The percentage of transmittance light radiation determines when light of certain frequency is passed through the samples. The spectrophotometer analysis records the intensity of absorption (A) or optical density (O.D) as a function of wavelength. The reaction mixture was taken in quartz cuvette and exposed to UV visible radition (200-800 nm) to MultiscanSky_1530-00496C spectrophotometer. The synthesized nanoparticles were well dispersed in the solution and were stable for long time by showing brown color to reaction mixture. The absorbance peak has already been recorded for nanoparticles which ranged from 2 to 100 nm in size. (Ravindra *et al*., 2014). The absorbance maxima was recorded at 421 nm which clearly indicated the formation of silver nanoparticles. (Fig.1)

**Fig. 1:**
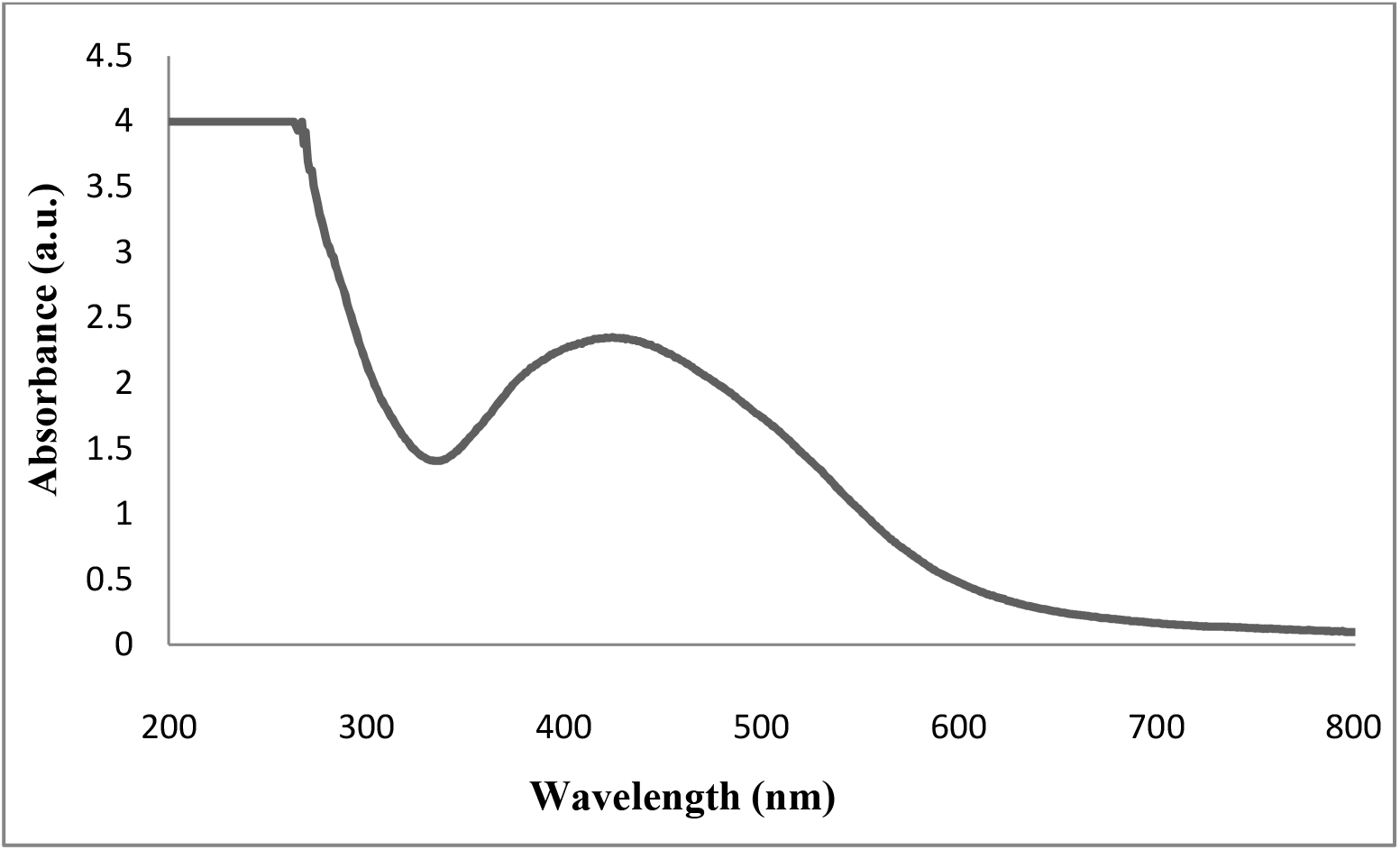
UV-vis absorption spectrum of Silver nanoparticles.

### SEM

The morphology and size of silver nanoparticles derived from *Drimia indica* leaf extract were characterized by Scanning Electron Microscope (Fig. 2). The image magnification was 10,000 X. From scanning electron microscope we obtained the average size of silver nanoparticles between 50-80 nm. The nanoparticles were spherical in shape and agglomerated. These synthesized nanoparticles were highly stable in nature.

**Fig. 2 :**
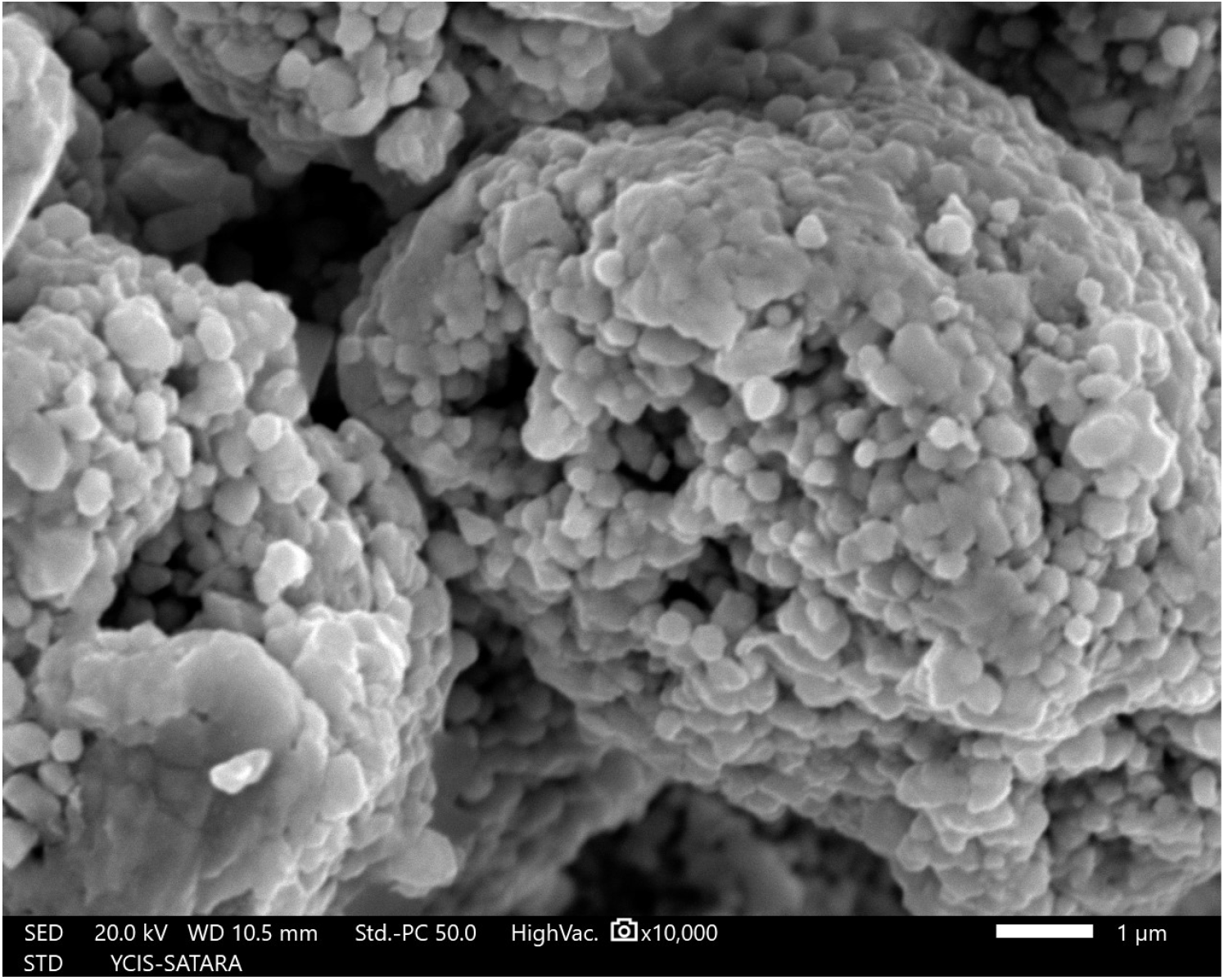
SEM micrograph of Silver nanoparticles.

### XRD

To investigate the crystalline nature and structure of Ag NPs, the sample was analyzed on BRUKER-D8 ADVANCE machine at CFC, Shivaji University, Kolhapur. The X-ray diffraction pattern obtained for the silver nanoparticles synthesized from *Drimia indica* leaf extract showed five distinct diffraction peaks (Fig. 3) of 2θ values of 38.11*°* (111), 44.30*°*(200), 64.22*°* (220), 77.51 *°* (311)and 81.47*°* (222) with the assigned planes indicated in the braktes. The free crystalline nature of synthesized nanoparticles was explained on the basis of JCPDS file. Analysis was done by using Origin Pro 8 software.

**Fig. 3 :**
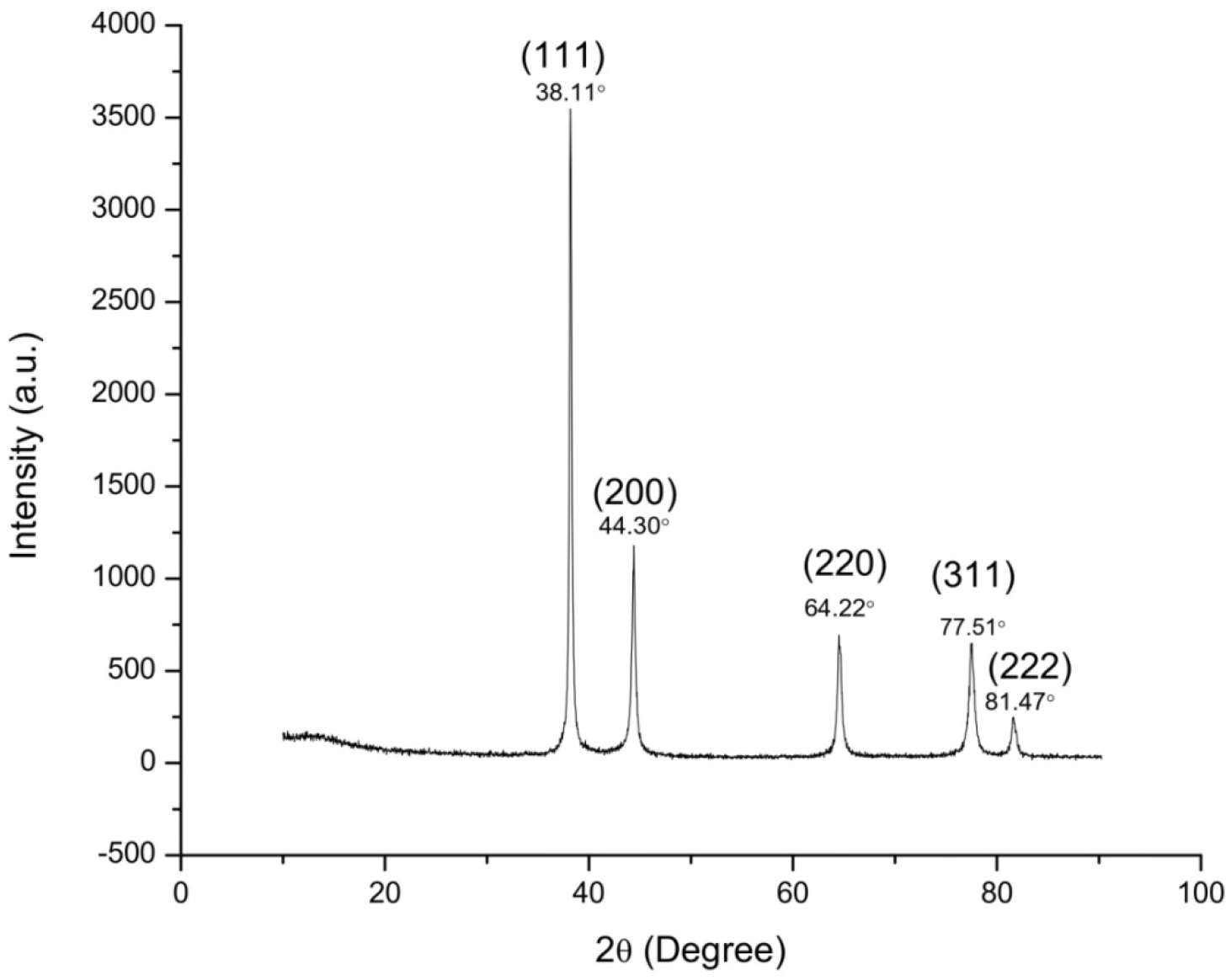
X-ray diffraction pattern of Silver nanoparticles.

### Antimicrobial assay

The antibacterial activity of silver nanoparticles synthesized from *Drimia indica* leaf extract was tested on pathogenic bacteria like *Pseudomonas aeruginosa* [ATCC2021], *Klebsiella pneumonia* [ATCC2075] and *Escherichia coli* [ATCC2065]. Bacterial cultures were maintained in liquid nutrient broth. Calculation of inhibition activity of synthesized nanoparticles on selected pathogenic bacterial cultures was done using different concentrations of Ag NPs. Antibacterial assay was carried out in 96-well microtiter plate. The assay volume was set to 300 µl which contained 260 µl nutrient broth, 20 µl bacterial culture and 20 µl Ag NPs. The Ag NPs were used in varied concentrations (25 µl/ml, 50 µl/ml and 100 µl/ml from mg/ml stock). Replacement of 20 µl Ag NPs with 20 µl distilled water in assay mixture served as negative control. The experiment was performed in triplicates. The 96-well microtitre plate was incubated for 12 hours. The readings were recorded after time interval of every 30 minutes in MultiscanSky_1530-00496C spectrophotometer. Graph was plotted which indicated sigmoid growth curve. Similar assay was repeated for all bacteria under study. The lowest concentration which inhibits the growth of bacterial strain is taken as MIC of the Ag NPs for the pathogenic bacteria. Broad spectrum antibiotic streptomycin in different concentrations (25 µl/ml, 50 µl/ml and 100 µl/ml) was used to calculate the MIC values for bacterial strains against streptomycin.

When Ag NPs were tested for antimicrobial activity, *Pseudomonas aeruginosa* showed MIC value of 25 µl/ml, *Klebsiella pneumoniae* showed 25 µl/ ml and *Escherichia coli* showed 50 µl/ ml (Table 1).

**Table. 1.** MIC Values of Silver nanoparticles with Streptomycin.

| Sr. No. | Bacteria | MIC values $\mu\text{l/ml}$ | MIC values $\mu\text{l/ml}$ |
| --- | --- | --- | --- |
|  |  | Ag NP's | Streptomycin |
| 1. | *Pseudomonas aeruginosa* | 25 | 100 |
| 2. | *Klebsiella pneumonia* | 25 | 50 |
| 3. | *Escherichia coli* | 50 | 100 |

*Pseudomonas aeruginosa* inhibition percentage (Fig. 4) was recorded to be highest (68%) at 100 µl/ml concentration of streptomycin while it was found to be lowest (45%) at 25 µl/ml Streptomycin concentration. The highest inhibition percentage (65%) at 100 µl/ml and lowest (43%) at 25 µl/ml was recorded for *Klebsiella* pneumoniae. While in case of *Escherichia coli* antibiotic Streptomycin showed highest inhibition percentage (69%) at 100 µl/ml and lowest inhibition percentage (47%) at 25 µl/ml (Fig. 4).

**Fig. 4.**
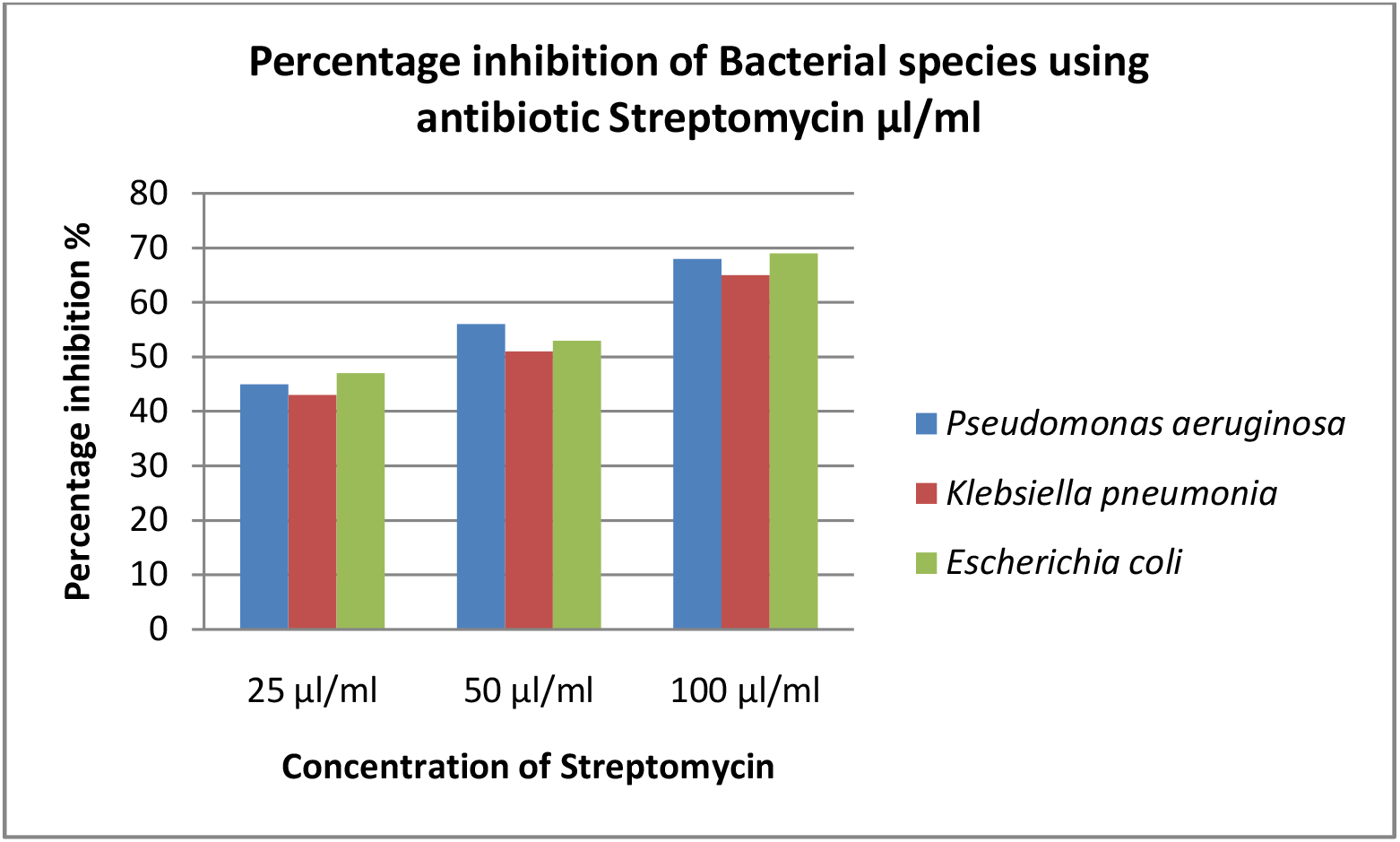
Percentage inhibition of Bacterial species using antibiotic Streptomycin.

The highest inhibition percentage of Ag NP’s against *Pseudomonas aeruginosa* (Fig.5) was observed to be 54% at Ag NPs concentration 100 µl/ml and lowest 39% at 25 µl/ml. In case of *Klebsiella pneumoniae* highest inhibition percentage (49%) was observed at 100 µl/ml of concentration Ag NP’s and lowest was 35% at 25 µl/ml. While, highest percentage inhibition for *Escherichia coli* was 51% at 100 µl/ml and lowest was 37% at 25 µl/ml concentration of Ag NPs (Fig. 5).

**Fig. 5.**
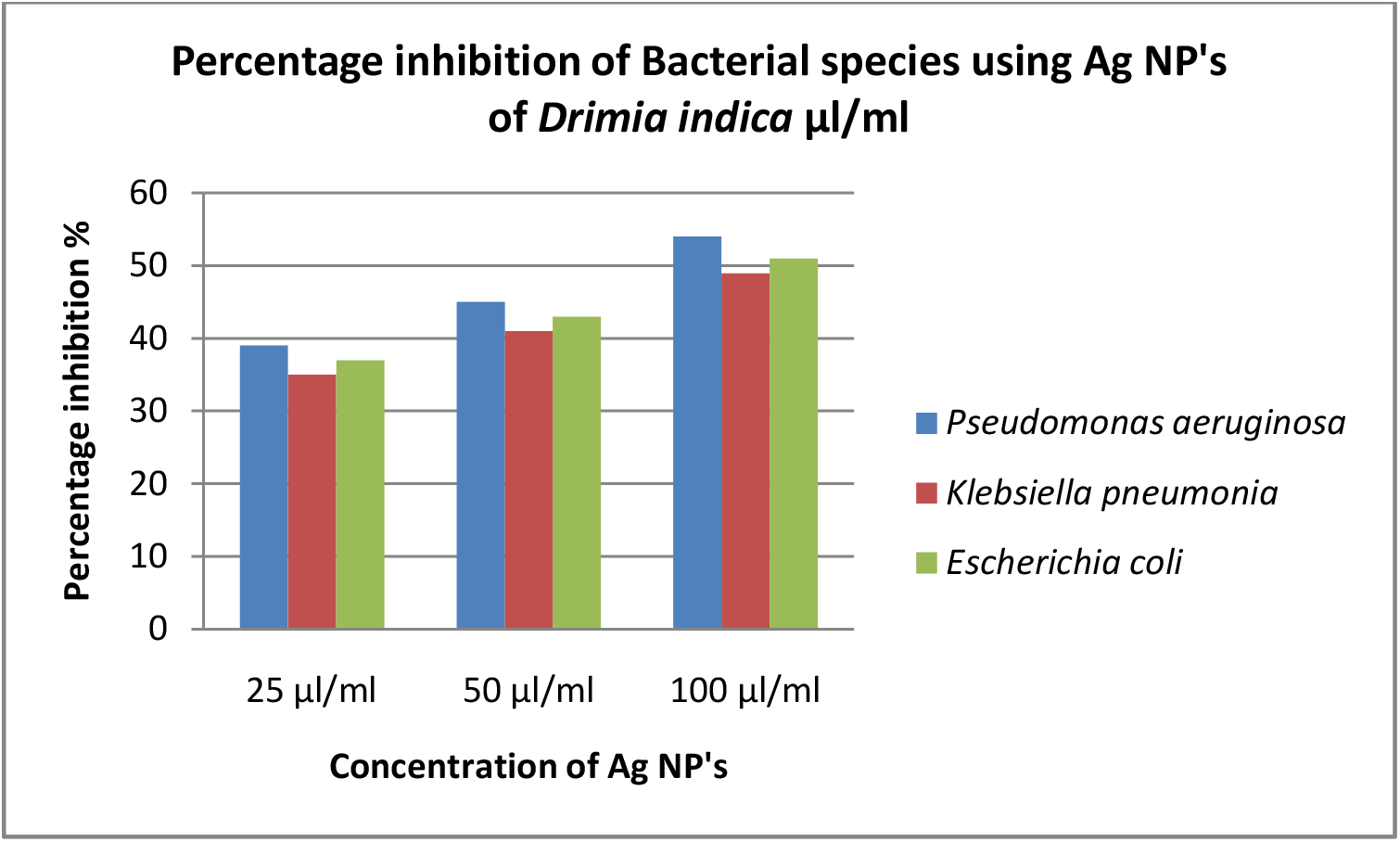
Percentage inhibition of Bacterial species using Ag NP’s of *Drimia indica*.

Thus, while studying the antibacterial activity of *Pseudomonas aeruginosa*, highest inhibition percentage (68%) was recorded when Streptomycin (100 µl/ml) was used and when Streptomycin was replaced with same concentration of Ag NPs, the highest inhibition percentage recorded was 54 %. In case of *Klebsiella pneumoniae*, highest inhibition percentage (65 %) was recorded in Streptomycin (100 µl/ml). The same concentration of Ag NPs exhibited 49 % inhibition. Highest inhibition percentage (69 %) was recorded in *Escherichia coli* for Streptomycin (100 µl/ml). Highest inhibition percentage 51 % was recorded for *Escherichia coli* when Ag NP’s (100 µl/ml) was used.

The results depicted that the role of antibiotic was remarkable in inhibition of growth in *Pseudomonas aeruginosa, Klebsiella pneumoniae* and *Escherichia coli*. Comparative results avoiding antibiotic and replacing it with biologically synthesized Ag NP’s showed the presence of inhibition activity in all the bacteria tested. Although inhibition activity of Ag NPs was lower than that of antibiotic, the Ag NPs were able to show antibacterial activity. Further research needs to be refined by designing the protocol in which higher concentrations must be tried to determine antibacterial activity. However, high doses of Ag NPs have advantage over higher doses of antibiotics as Ag NPs are ecofriendly, nontoxic and biologically synthesized. Biologically synthesized nanoparticles can be used as better options over hazardous antibiotics by using them as antibacterial agents.

## Conclusion

Ag NPs of *Drimia indica* has shown the lowest antibacterial effect than control in present study, but when the concentration were increased antibacterial effect also increased. It clearly revealed that the inhibition activity of Ag NPs is completely dose dependent and it inhibits the pathogenic bacteria. Thus, they can be widely used as medicines for bacterial diseases. This will eventually help to investigate the problems and mutations caused by high doses of antibiotics.

## Notes

### Competing Interest Statement

The authors have declared no competing interest.

### Summary of Updates

The file was yet not uploaded. I think the process done was half the way by mistake. Herewith doing the complete submission now

